# Effect of age and sex on gene expression-based radiation biodosimetry using mouse peripheral blood

**DOI:** 10.1101/2022.10.27.514053

**Authors:** Constantinos G. Broustas, Igor Shuryak, Axel J. Duval, Sally A. Amundson

**Affiliations:** Center for Radiological Research, Columbia University Vagelos College of Physicians and Surgeons, Columbia University Irving Medical Center, New York, NY, USA

**Keywords:** age, sex, radiation, gene expression, mice

## Abstract

Blood-based gene expression profiles that can reconstruct radiation exposure are being developed as a practical approach to radiation biodosimetry. However, age and sex could potentially limit the accuracy of the approach. In this study, we determined the impact of age on the peripheral blood cell gene expression profile of female mice exposed to radiation and identified differences and similarities with a previously obtained transcriptomic signature of male mice. Young (2 months) and old (24 months) female mice were irradiated with 4 Gy X-rays, total RNA was isolated from blood 24hr later and subjected to whole genome microarray analysis. Dose reconstruction analyses using a gene signature trained on gene expression data from irradiated young male mice showed accurate reconstruction of 0 or 4 Gy doses with root mean square error of ± 0.75 Gy (R^2 = 0.90) in young female mice. Although dose reconstruction for irradiated old female mice was less accurate than young female mice, the deviation from the actual radiation dose was not statistically significant. Pathway analysis of differentially expressed genes revealed that after irradiation, apoptosis-related functions were overrepresented, whereas functions related to quantities of various immune cell subtypes were underrepresented, among differentially expressed genes from young female mice, but not older animals. Furthermore, young mice significantly upregulated genes involved in phagocytosis, a process that eliminates apoptotic cells and preserves tissue homeostasis. Both functions were also overrepresented in young, but not old, male mice following 4 Gy X-irradiation. Lastly, functions associated with neutrophil activation that is essential for killing invading pathogens and regulating the inflammatory response were predicted to be uniquely enriched in young but not old female mice. This work supports the concept that peripheral blood gene expression profiles can be identified in mice that accurately predict physical radiation dose exposure irrespective of age and sex. However, inclusion of age and sex as biological factors is essential for effectively predicting radiation injury and for developing radiation medical countermeasures.

## Introduction

Early blood-based gene expression signatures that can predict radiation dose exposure are being developed as a practical approach to radiation biodosimetry for use in a large-scale radiological event, such as an intentional detonation of an improvised nuclear device or nuclear accident [Amundson, 2022]. Moreover, pathway analysis of differentially expressed genes could provide valuable biological information that could be predictive of later damage, thus, guiding the implementation of appropriate long-term medical management strategies, or enabling the development of novel medical countermeasures. However, the development of accurate and rapid biodosimetry tools represents a significant challenge because several biological confounding factors, such as sex, age, and various comorbidities, such as pre-existing infections, could potentially limit the accuracy of blood-based gene expression signatures of radiation exposure and, thus, useful radiation biomarker candidates should be accurate, at least, across age and sex.

Individuals exposed to radiation show different degrees of sensitivity depending on their age. It has been shown that sensitivity to radiation is high at early ages, declines during middle life, and again increases in older adults [Hernandez et al., 2015]. Similar trends are also found in mice [Hernandez et al., 2015]. Increased reactive oxygen generation with a concomitant decline in the activity of antioxidant enzymes, telomere attrition, and impaired DNA damage response and repair are some of the factors that contribute to the altered sensitivity to radiation in aged individuals [Hernandez et al., 2015]. Furthermore, molecular analysis of human blood samples revealed significant age-related changes in gene expression [Peters et al., 2015] that lead to dysregulation of immune function that involves both the adaptive and the innate immune system. These changes result in an aberrant chronic low-grade pro-inflammatory state [Goronzy et al., 2013; Fulop et al., 2018], believed to occur to a greater extent in females than males [Furman et al., 2014].

In addition to age, sex is a major determinant of variation in physiology and disease susceptibility in humans and many immunological and inflammatory diseases have a striking gender bias in incidence and severity [Klein and Flanagan, 2016], such as in the immunity and clinical manifestations of infectious or autoimmune diseases and malignancy [Klein and Flanagan, 2016]. Immune systems of men and women function and react to infections and vaccination differently [Giefing-Kröll et al., 2015] and respond differently to immune cell stimuli, such as cytokines and microbes [Bakker et al., 2018; Piasecka et al., 2018]. The number and activity of innate immune cells, including macrophages and dendritic cells, as well as inflammatory immune responses, are higher in females than males [Boissier et al., 2003; Xia et al., 2009; Melgert et al., 2010]. For some infectious diseases, an inability to properly clear or control a pathogen may contribute to increased prevalence and severity of disease in males as compared with females, and epidemiological studies have shown that males have a higher mortality rate to various infectious diseases [vom Steeg and Klein, 2016]. However, heightened inflammatory, and cellular immune responses in females, though essential for pathogen clearance and increased vaccine efficacy, may underlie increased development of symptoms of disease among females as compared with males following infection.

In humans, the impact of age and sex on immune phenotypes has been described [Marquez et al., 2020] and sex-dependent variation in white blood cell gene expression in healthy human populations has been reported [Whitney et al., 2003] that could potentially limit the accuracy of blood-based gene expression radiation biodosimetry. However, the accuracy of radiation biodosimetry appears less affected by sex in human subjects [Paul and Amundson, 2011], whereas, in a murine radiation biodosimetry study, it was shown that sex differences had an impact on gene expression signatures following radiation and influenced the accuracy of the prediction of radiation exposure at low doses (0.5 Gy), albeit not at higher doses (1 and 2 Gy) [Meadows et al., 2008]. Thus, sex-related differences could be a concern for distinguishing low dose exposures from unirradiated controls. On the other hand, we demonstrated that age modulated the transcriptomic profile of blood cells in male mice exposed to radiation, and that numerous biological processes and pathways were altered, while it compromised the accuracy of prediction of physical radiation dose exposure [Broustas et al., 2021].

In this study, we examined the impact of age on white blood cell gene expression profiles in response to acute radiation exposure using female C57BL/6 mice and compared the data with those from male mice treated under the same conditions [Broustas et al., 2021]. The transcriptomic response was analyzed 24 h following 4 Gy X-irradiation to match early monitoring applications, whereas whole blood was chosen as a minimally-invasive source for the development of molecular biomarkers. We applied a recently developed radiation dose reconstruction method [Ghandhi et al., 2019] to determine the impact of age and sex on this method of radiation biodosimetry. Furthermore, we performed gene ontology analysis to identify and compare differentially enriched pathways, diseases, and functions in mice of different age and sex.

## Materials and Methods

### Animals and irradiation

A total of 20 female C57BL/6 mice were purchased from Jackson Labs (10 young [2 months] and 10 old [24 months] at the time of irradiation) and acclimated for 2 weeks before irradiation. Each treatment group consisted of 5 mice and the treatment was as follows: (1) Control (unirradiated)-young mice, (2) Control (unirradiated)-old mice, (3) 4 Gy X-ray-young mice, (4) 4 Gy X-ray-old mice. Mice were fed the standard chow, without caloric restriction. All animal experiments were conducted in accordance with applicable federal and state guidelines based on approved by the Columbia University Institutional Animal Care and Use Committee (approval number AC-AAAT6450). Mice were either sham-irradiated or exposed to 4 Gy X-rays of total body irradiation from an X-RAD 320 Biological Irradiator (operating at 320kV, 12.5mA with a 2 mm Al filter [HVL∼1.0 mmCu]) at a dose rate of 1 Gy/min. The male study has been published previously [Broustas et al., 2021] and performed under the same conditions described herein.

### Blood Collection and RNA isolation

Blood was collected 1 day post-irradiation by cardiac puncture at the time of euthanasia (by CO_2_ asphyxiation). Each sample (∼0.4 ml blood) was added to a 15 ml centrifuge tube that contained 1.6 ml of PAXgene Blood RNA stabilization and lysis solution (PreAnalytix GmBH) and mixed thoroughly, while a small amount of blood was added to sodium EDTA anti-coagulant containing tubes for blood count using a Genesis hematology system (Oxford Science). After collection, blood was incubated at 4 °C for 24 h. RNA was purified following the PAXgene RNA kit recommendations with on-column DNase I treatment. As excessive amounts of globin transcript have been shown to interfere with gene expression signatures derived from blood, globin RNA was reduced using the Ambion GLOBINclear-mouse/rat kit (Thermofisher). RNA yields were quantified using the NanoDrop ND1000 spectrophotometer (Thermofisher) and RNA quality was checked by the 2100 Bioanalyzer (Agilent). High quality RNA with an RNA integrity number of at least 7.0 was used for microarray hybridization.

### Microarray hybridization

Cyanine-3 labeled cRNA was prepared using the One-Color Low input Quick Amp Labeling kit (Agilent). Dye incorporation and cRNA yield was measured with a NanoDrop ND1000 spectrophotometer (Thermofisher). Labeled cRNA was fragmented and hybridized to Agilent Mouse Gene Expression 4×44K v2 Microarray Kit (G4846A). Slides were scanned with the Agilent DNA microarray scanner (G2505B) and the images were analyzed with Feature Extraction software (Agilent) using default parameters for background correction and flagging non-uniform features.

### Data analysis

Background-corrected hybridization intensities were imported into BRB-ArrayTools, version 4.5.1 [Wright and Simon, 2003], log_2_-transformed and median normalized. Non-uniform outliers or features not significantly above background intensity in 25% or more of the hybridizations were filtered out. In addition, a minimum 1.5-fold change in at least 20% of the hybridizations was set as a requirement. Furthermore, probes were averaged to one probe per gene and duplicate features were reduced by selecting the one with maximum signal intensity. Class comparison was conducted in BRB-ArrayTools to identify genes differentially expressed between radiation exposed samples and matched unirradiated controls using a random variance t-test. Genes with p-values less than 0.005 were considered statistically significant. The false discovery rate (FDR) was estimated for each gene by the method of Benjamini and Hochberg [Hochberg and Benjamini, 1990], to control for false positives. The cutoff in this analysis was set at an FDR of less than 0.05. Hierarchical clustering of microarray gene expression data was performed with the Dynamic Heatmap Viewer of the BRB-ArrayTools software using a one minus correlation metric and average linkage. Genes differentially expressed following exposure to 4 Gy X-ray at day 1 in the young female mice from this study, or in the young male mice [Broustas et al., 2021] were used to construct the heatmap. Venn diagrams [Bardou et al., 2014] were used to identify unique and overlapping differentially expressed genes from irradiated old and young female mice.

### Dose reconstruction

We analyzed 4 data sets of gene expression in mice: young males, old males, young females, and old females. The young male data set was a priori assigned as a “training” data set for identifying radiation-responsive gene candidates, whereas the other 3 data sets were assigned as “testing” sets where these candidate genes would be applied for dose reconstruction. The search for candidate genes in the “training” young male data was performed by selecting those genes which had a statistically significant positive or negative Pearson correlation coefficient with dose (either 0 or 4 Gy) and were measured (not missing) in all 12 samples. A p-value threshold of 0.05 with Bonferroni correction for multiple comparisons was selected to indicate statistical significance.

Top 10 candidate upregulated genes and top 10 downregulated genes from the young males “training” data set (i.e., genes with positive or negative statistically significant Pearson correlation coefficients with radiation dose) were selected for further analysis. Their signals were combined using geometric means, separately for upregulated and downregulated groups. These top 10 upregulated and downregulated genes, originally identified in the young males, were then selected in each of the three “testing” data sets: old males, young females and old females. The geometric means of top 10 upregulated and downregulated genes were calculated for all 4 data sets.

The net signal was defined as the difference in geometric means between upregulated and downregulated genes. A linear regression model was fitted to the “training” young male data. The dependent (target) variable was radiation dose, and the independent (predictor) variable was the net signal of the genes. An intercept term was also allowed. The best-fit parameters for the regression model obtained on “training” data were then used to make dose predictions (reconstructions) on the 3 “testing” data sets. Root mean squared errors (RMSE) and coefficient of determination (R^2^) performance metrics, which compared actual with reconstructed dose values, were calculated to evaluate the model. Potential differences in dose reconstructions as function of sex (male or female) and age (young or old) were investigated by ANOVA and Tukey post-hoc test. These analyses were performed in R 4.2.0 software.

### Gene ontology analysis

Lists of genes that were either significantly overexpressed or underexpressed compared with controls were analyzed using the Ingenuity Pathway Analysis (IPA) core pathway (Qiagen Ingenuity Systems) to identify significantly affected canonical pathways, diseases and functions. Benjamini corrected p values of < 0.05 were considered significant.

## Results

### Effect of radiation on blood cell counts of young and old mice

We analyzed the differential blood cell counts from young (2 mo) and old (24 mo) female mice that had been exposed or not to 4 Gy X-ray irradiation. Overall, the number of white blood cells (WBC) in older mice was significantly higher than in younger mice (p = 7.21E-03) (Fig. 1a). However, the total number of WBC 24 h post-irradiation was similar in both age groups (p = 4.95E-01). The relative percentage of lymphocytes in unirradiated female mice was 60-70% of the total WBC number and similar between young and old animals (p = 6.84E-02). Following radiation, relative lymphocyte percentages fell to approximately 35% that were significant compared with unirradiated animals (p = 1.33E-04 for young and p = 1.32E-02 for old mice). In contrast, the relative percentage between young and old mice following irradiation were similar (p = 2.20E-01). These results were similar in male mice exposed to radiation under the same conditions [Broustas et al., 2021]. Furthermore, neutrophils, which are less sensitive to radiation than lymphocytes, became the most abundant species after irradiation, accounting for approximately 50-70% of the WBC (Fig. 1b). A significant increase in the percentage of neutrophils was found in irradiated young (p = 3.74E-04) and old (p = 3.10E-03) mice compared with unirradiated animals (Fig. 1c). There was also a statistically significant difference in the percentage of neutrophils from young and old mice before irradiation (p = 4.12E-02) and the difference remained significant after irradiation (p = 2.58E-02). The percentage differentials between irradiated young female and male mice were not significant (p = 2.47E-01), whereas between irradiated old female and male mice were marginally significant (p = 4.99E-02). In comparison, geriatric male mice have higher percentages of neutrophils at steady state (p = 3.89E-02), but unlike female mice, there was no statistical difference after irradiation (p = 7.15E-02) [Broustas et al., 2021]. Moreover, percentages of monocytes, which differentiate into macrophages and dendritic cells, were higher in older female mice (p = 2.47E-02) under basal conditions compared with young female mice; however, exposure to 4 Gy X-rays did not appreciably change the relative abundance in young (p = 2.92E-01) or older (p = 7.43E-02) female mice (Fig. 1d), as seen with male mice [Broustas et al., 2021]. In contrast to male mice [Broustas et al., 2021], platelet counts were not significantly elevated in older mice compared with young mice (p = 1.21E-01) under basal conditions. Radiation exposure did not change platelet counts in either young (p = 2.02E-01) or old mice (p = 2.81E-01) compared with unirradiated mice. However, the number of platelets was higher in irradiated old mice compared with irradiated young animals (p = 9.71E-03), suggesting that platelets from older mice are more resistant to radiation or that platelets from older mice respond to radiation with a delayed kinetics. Comparing male and female platelets counts, baseline counts were similar (p = 4.15E-01) between young and old female and male mice; however, older male mice had a significantly higher cell count than older female mice (p = 6.36E-03). Finally, exposure to irradiation preserved platelets from young male mice better than young female mice (p = 3.14E-03), while there was no such protection between old mice (p = 1.85E-01).

**Figure 1.**
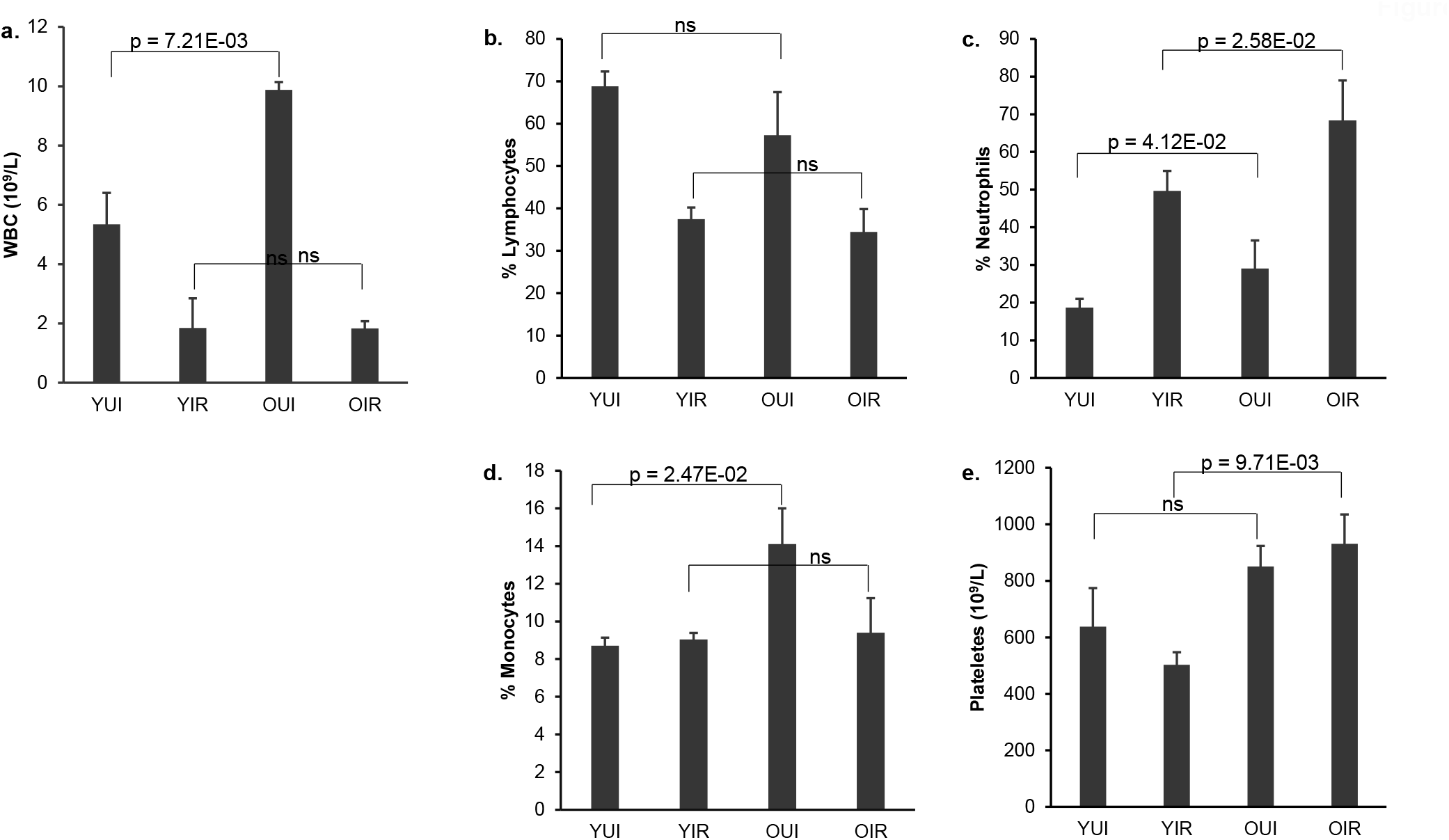
Total white blood cell count (WBC) and percentages of different blood types in young and old female mice 1 day after 4 Gy X-irradiation. (a) total WBC counts, (b) lymphocytes (%), (c) neutrophils (%), (d) monocytes (%), (e) platelets counts. Data represent the mean S.E.M (n = 3). P values were calculated using the unpaired Student’s t-test. ns: not significant (p ≥ 0.05). YUI, young unirradiated; YIR, young irradiated; OUI, old unirradiated; OIR, old irradiated.

### Microarray Analysis

Global gene expression was measured in the blood of young and old C57BL/6J female mice using Agilent’s whole mouse genome microarrays. Class comparison using BRB-ArrayTools [Wright and Simon, 2003] identified a total of 2,505 and 2,242 differentially expressed genes (p < 0.005, false discovery rate (FDR) < 5%) in young irradiated vs. young control mice and old irradiated vs. old control mice, respectively (Fig. 2a; the complete list of differentially expressed genes can be found in online suppl. File 1). In a deviation from the previous studies with male mice [Broustas et al., 2017, 2021] that showed that the majority of the differentially expressed genes are downregulated in response to irradiation, 60% of the differentially expressed genes in the young female mice were upregulated, whereas 40% were downregulated. In the old female mice, 62% of the differentially expressed genes were upregulated and 38% were downregulated. At steady state, 2,352 genes were found to be differentially expressed in control (unirradiated) young female versus control aged female mice with 64% of the genes being downregulated in the geriatric population (Fig. 2a). In male mice, we found that 85% of the genes differentially expressed between young and old animals were downregulated in the older mice [Broustas et al., 2021]. A total of 3,919 genes were responsive to ionizing radiation in at least one age group. Of these genes, 828 were common between young and old female mice, whereas 1,677 genes (42.8%) in the “young” cohort and 1,414 genes (36.1%) in the “old” cohort were unique to the respective age group (Fig. 2b). Comparison of radiation-responsive genes between young male [Broustas et al., 2021] and young female mice revealed that 1,264 genes were common between the two genelists, which represented 26.4% of the differentially expressed genes in the young animals, whereas, there were only 671 common between old male and female or 14.3% of radiation responsive differentially expressed genes (Fig 2b and online suppl. Fig. S1).

**Figure 2.**
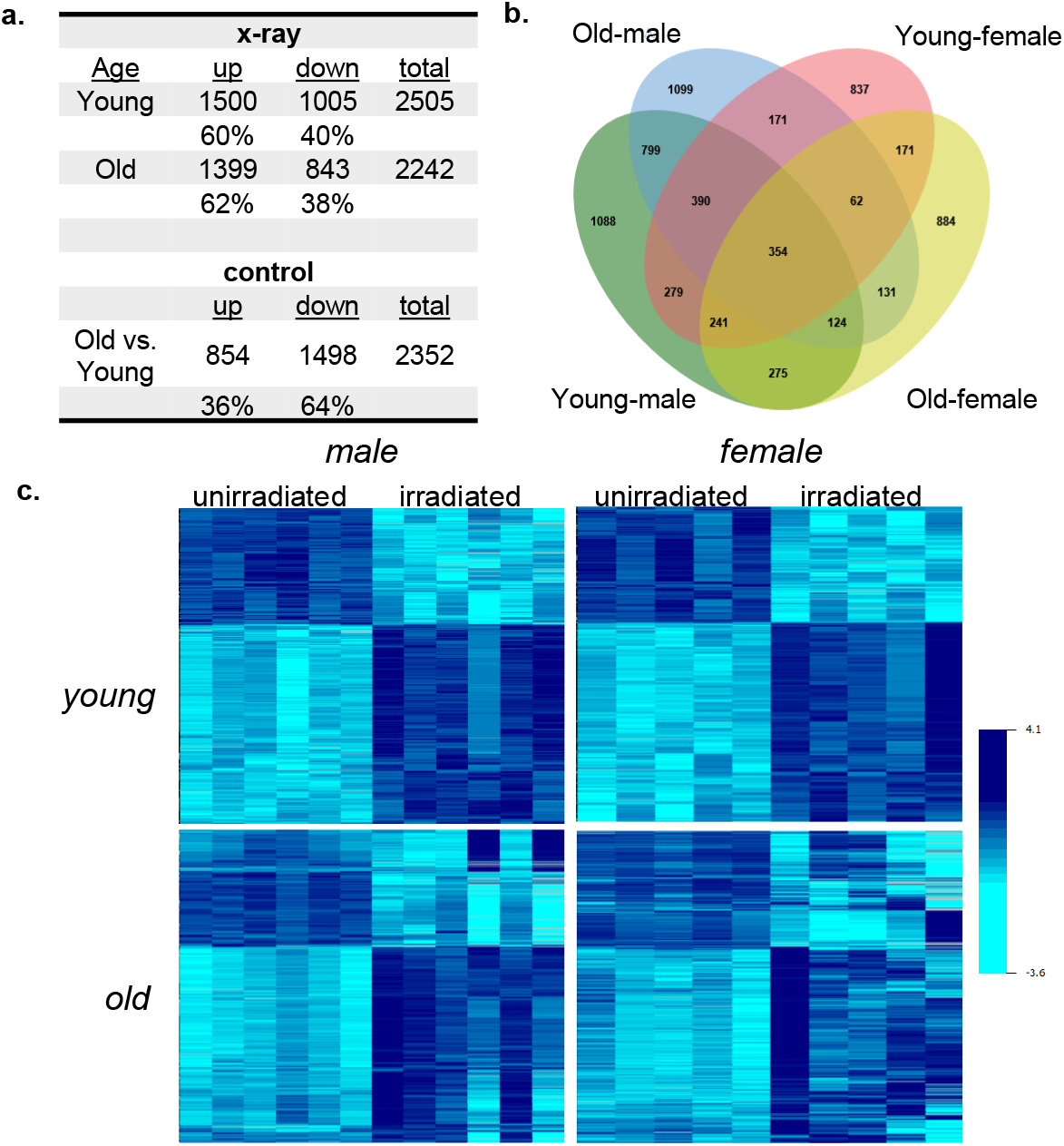
Differentially expressed genes. (**a**) Significantly differentially expressed genes in young and old female mouse blood (p < 0.005) following 4 Gy X-rays (X-ray) relative to unirradiated mice and under basal conditions comparing old versus young female mice (control). Percent of upregulated (up) and downregulated (down) genes are shown. (**b**) Venn diagram showing overlap of differentially expressed genes in irradiated young male and female versus irradiated old male and female mice (**c**) Heatmaps of combined effects of age and sex on commonly differentially expressed genes in young female and young male mice exposed to radiation.

To determine the effect of age on gene expression in response to radiation, we followed the expression of the 1,264 radiation-responsive genes that are common between young male and female mice and assessed their expression levels in aged mice from both sexes. Heatmap generation revealed that expression of these genes in response to irradiation diverged significantly among replicates of geriatric mice, especially in the older female mice (Fig. 2c).

### Impact of age and sex on reconstruction of radiation exposure

To assess the impact of age and sex on radiation-response gene signatures, we identified 354 genes that were commonly differentially expressed in response to radiation irrespective of age or sex (Fig. 2b) and compared fold-change gene expression using the young male fold-changes as reference values. Plotting fold changes in response to irradiation against the fold change of male young mice revealed that the fold-change values from either old male or old female mice differed dramatically from those of young male or female irradiated mice (p = 1.82E-04 young *vs*. old male; p = 3.82E-13 young *vs*. old female mice) (Fig. 3) with geriatric mice showing markedly lower fold-changes than those of young animals. Comparison of sex differences showed that young female fold-changes were very similar to those of young male mice (p = 7.83E-02). In contrast, there was a statistical difference in fold-changes between aged female and aged male mice (p = 1.42E-29).

**Figure 3.**
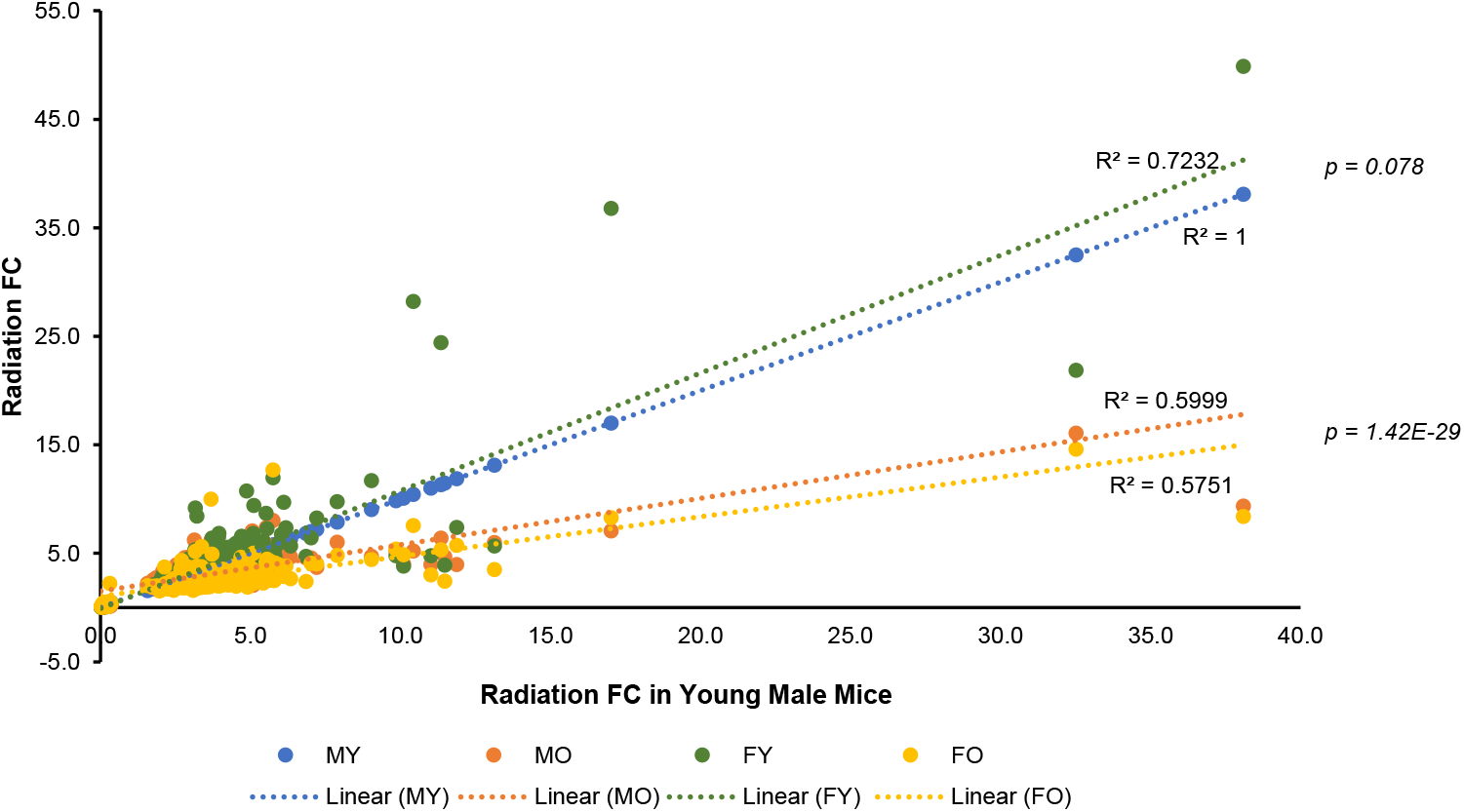
Impact of age and sex on the fold-change of radiation response signature genes. Fold changes (FC) in differentially expressed genes from irradiated mice of different age and sex compared with radiation fold changes (FC) of differentially expressed genes from irradiated young male mice. R^2^ values represent the fit of the gene fold changes to a line. P values were calculated using the paired Student’s t-test. MY: male young; MO: male old; FY: female young; FO: female old.

Next, we attempted a dose reconstruction analysis using the transcriptomic profile of young male mice [Broustas et al., 2021] as the training set. The initial search for genes significantly correlated with radiation dose in the “training” young male data identified 189 genes. The top 10 downregulated and top 10 upregulated genes from this group are shown in Table 1. Most of these genes showed similar dose-responsive trends in the “testing” data sets (*i*.*e*., signs and magnitudes of Pearson correlation coefficients were similar to those in the “training” data), although they did not always reach statistical significance. The best-fit parameters for a linear regression which predicted radiation dose on the “training” young male data were: intercept = 1.083 ± 0.073 (standard error), p-value = 4.00E-08; slope (dependence on gene net signal) = 0.552 ± 0.018, p-value = 4.10E-11. Although the lists of significantly dose-correlated genes were different for different ages/sexes, those genes that were strongly upregulated or downregulated with dose (top 10 candidates) identified in the “training” young males data showed similar dose response trends in the other ages/sexes, even though these trends did not always reach statistical significance there. So, using geometric means of these 10 upregulated and downregulated genes allowed decently accurate dose reconstruction in the “testing” data sets (old males, young and old females). ANOVA and Tukey post-hoc test analyses suggested that reconstructed doses did not significantly vary with sex (adjusted p-value = 0.80) or age (adjusted p-value = 0.67). A table comparing actual with reconstructed doses is shown in Table 2. Overall, the root mean square error (RMSE) on testing data was 0.75 (R^2^ = 0.90).

**Table 1.** Top 10 downregulated and top 10 upregulated dose-responsive genes in young male data.

| Symbol | Correlation coefficient with dose | Bonferroni adjusted p-value |
| --- | --- | --- |
| Fcmr | -0.991 | 6.65E-06 |
| Hvcn1 | -0.990 | 1.18E-05 |
| Pla2g2d | -0.994 | 1.43E-05 |
| H2-Oa | -0.988 | 3.64E-05 |
| H2-Ob | -0.988 | 3.85E-05 |
| Tnfrsf13b | -0.987 | 4.46E-05 |
| Ighd | -0.987 | 4.62E-05 |
| Btla | -0.987 | 5.00E-05 |
| Ccr7 | -0.987 | 5.85E-05 |
| Ccr6 | -0.986 | 7.05E-05 |
| Lgals3bp | 0.984 | 1.38E-04 |
| Ifi27l2a | 0.982 | 2.36E-04 |
| Phlda3 | 0.982 | 2.66E-04 |
| Oas2 | 0.979 | 5.85E-04 |
| Oasl2 | 0.973 | 1.76E-03 |
| Gabarapl1 | 0.973 | 1.99E-03 |
| Cdkn1a | 0.971 | 2.43E-03 |
| Oas1a | 0.967 | 4.84E-03 |
| Lamp1 | 0.967 | 5.07E-03 |
| Rtp4 | 0.967 | 5.20E-03 |

**Table 2.** Comparison of actual and reconstructed radiation doses.

| Actual vs reconstructed doses |  |  |  |  |
| --- | --- | --- | --- | --- |
| Sex | Age | Actual dose (Gy) | Reconstructed dose (Gy) |  |
|  |  |  | *mean* | *sd* |
| M | Young | 0 | 0.022 | 0.11 |
|  | Young | 4 | 3.978 | 0.30 |
|  | Old | 0 | 0.363 | 0.17 |
|  | Old | 4 | 3.566 | 0.45 |
| F | Young | 0 | 0.424 | 0.70 |
|  | Young | 4 | 3.523 | 0.32 |
|  | Old | 0 | 0.775 | 0.56 |
|  | Old | 4 | 2.949 | 0.35 |
| RMSE (Gy) on testing data |  |  | R^2 on testing data |  |
| 0.75 |  |  | 0.90 |  |

### Pathway Analysis

We used Ingenuity Pathway Analysis (IPA) to identify the most significantly enriched canonical pathways and diseases or functions among differentially expressed genes considering a Benjamini-corrected *p*-value of less than 0.05 to be significant. Moreover, the activation state of each process was determined by its *z*-score. A z-score of at least 2 indicated activation, whereas a z-score of -2 or less indicated inactivation/inhibition.

Of the 69 canonical pathways with a |z| ≥ 2, 48 or 69.5% were common to both sex groups and ages (online suppl. File 2). Nineteen (19) pathways were unique to young, irradiated male and female mice, whereas older mice showed a greater dissimilarity with only 2 (3%) pathways shared between male and female. IPA analysis revealed that 98 diseases and functions were predicted to be differentially regulated in irradiated mice. Of these, 33 (34%) functions were common to all age and sex groups, 60 (61%) were unique to young animals, whereas 5 (5%) were enriched specifically in older mice (Supplementary File 3). As we have seen previously using male mice [Broustas et al., 2021], immune cell apoptosis-related functions were overrepresented specifically in the younger female mice, but not in older mice (Fig. 4a and online suppl. Table S1a for actual z-scores). Functions related to B and T lymphocyte death were overrepresented, whereas quantity of B cells and antibody production were underrepresented in young female mice exposed to radiation, but not old female mice. Similar results were noted using male mice, as well (Fig. 4b and Table S1b) [Broustas et al., 2021]. Furthermore, another significant class of disease and functions that was overrepresented in young female animals was related to phagocytosis (Fig. 5a and online suppl. Table S2a for actual z-scores). A smaller number of phagocytosis-associated functions was also predicted to be overrepresented in older female mice (Fig. 5a). Furthermore, phagocytosis-related biofunctions were predicted to be associated with irradiated young animals of both sexes, whereas they were not differentially represented in old animals (Fig. 5b and online suppl. Table S2b for actual z-scores).

**Figure 4.**
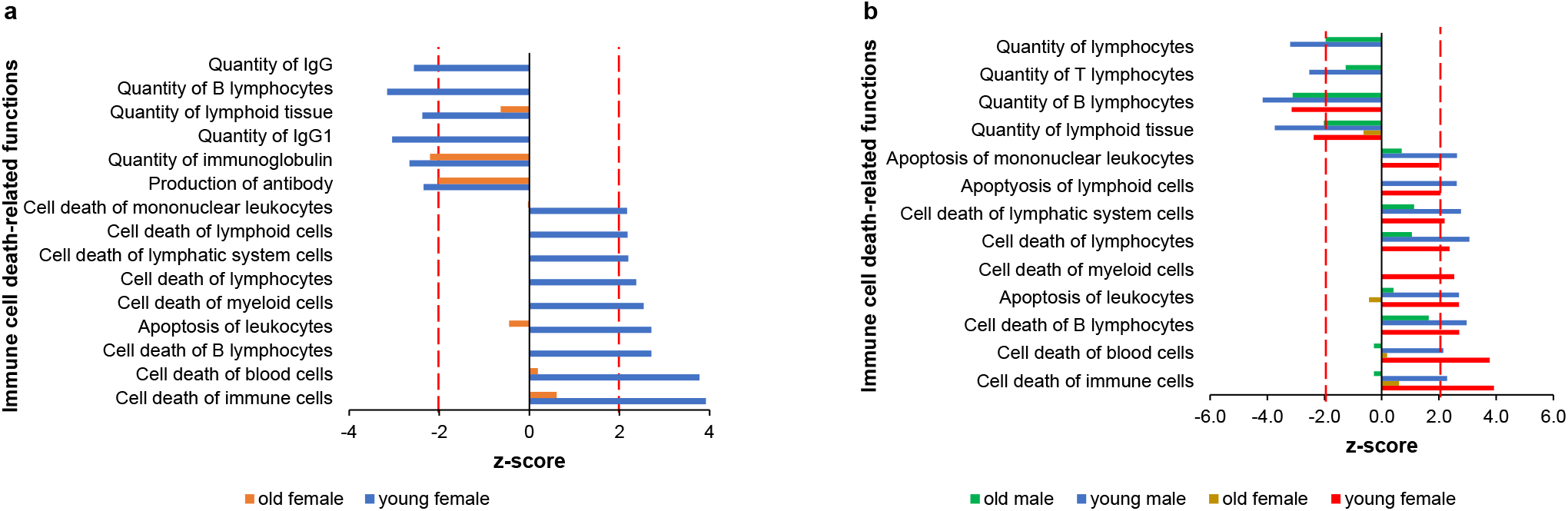
Immune cell death-related diseases and functions are overrepresented in irradiated young female mice. Significantly enriched cell death-related functions in young-irradiated versus old-irradiated female mice. Dotted line marks absolute z-score = 2.0.

**Figure 5.**
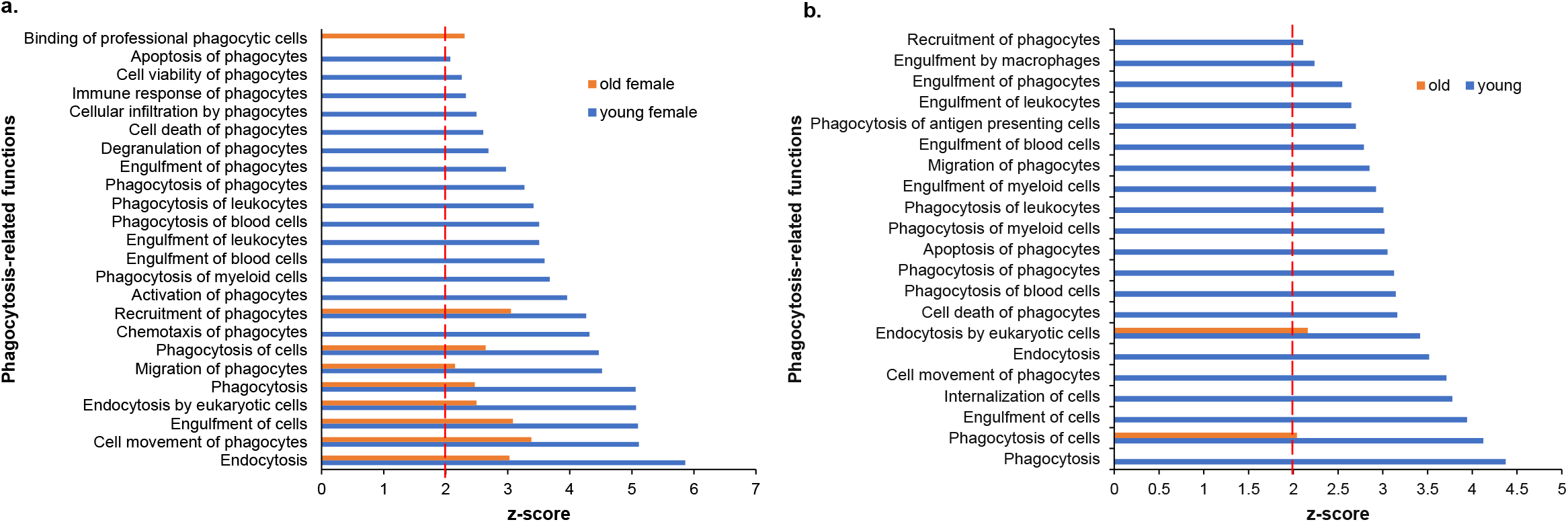
Phagocytosis-associated biological functions are overrepresented in irradiated young mice. (**a**) Phagocytosis-related functions significantly enriched in young-irradiated female mice versus old-irradiated female mice. (**b**) Phagocytosis-related functions overrepresented in young male and female mice versus old male and female animals after radiation exposure. Dotted red line marks z-score = 2.0.

Neutrophils, and other cells of the innate immune system, represent a first line of defense against pathogens and play a fundamental role in the activation, regulation, and orientation of the adaptive immune response. A unique feature of irradiated female mice was the overrepresentation of functions related to neutrophil recruitment and activation, which were significantly overrepresented only in young female mice (Fig. 6a and online suppl. Table S3a for actual z-scores values). Older female and young male, but not old male mice exposed to radiation were enriched in only one neutrophil-related function. Concomitantly, production and metabolism of reactive oxygen species were also predicted to be significant in young female and young male mice, but not older male animals, whereas differentially expressed genes from irradiated old female mice were enriched in these functions, albeit with significant lower z-scores (Fig. 6b and online suppl. Table S3b for actual z-scores values). Extending the analysis to myeloid cell related functions, in general, we found that myeloid-associated functions were overrepresented in young female and young male mice, less in older female mice, but were absent from older male mice following radiation (online suppl. Table S3c).

**Figure 6.**
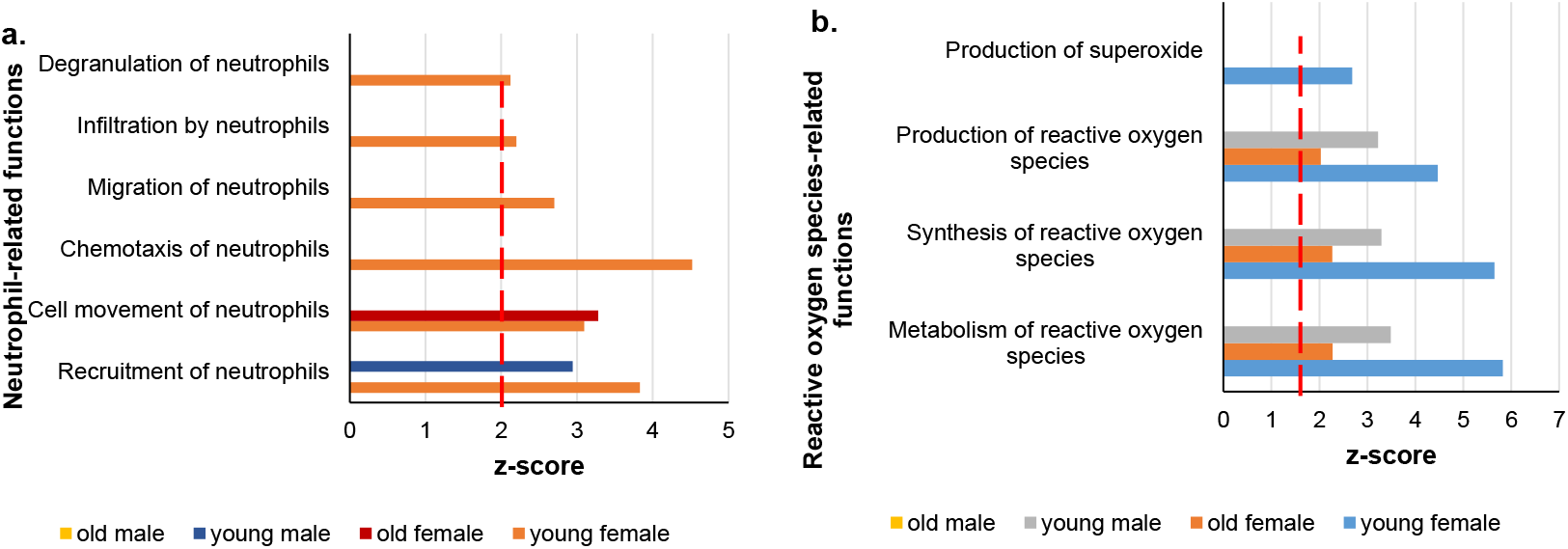
Neutrophil-related functions are enriched in young female mice exposed to radiation. (**a**) Significantly overrepresented neutrophil recruitment and activation functions. (**b**) Reactive oxygen generation and metabolism-related pathways in irradiated mice of different age and sex. Dotted red line marks z-score = 2.0

## Discussion

The purpose of radiation biodosimetry is to reconstruct radiation dose, as a surrogate of radiological injury. Large-scale transcriptomic analysis using whole blood has been used to select genes that may detect radiation exposure both in humans and mice [Dressman et al., 2007; Paul and Amundson, 2011; Broustas et al., 2017a, 2017b, 2018; Mukherjee et al., 2019a; Li et al., 2019]. Furthermore, gene ontology analysis of the differentially expressed genes has helped to identify critical biological functions that could be used to rationally design new mitigators against the deleterious effects of radiation. However, both physical factors, such as radiation quality, and biological factors, such as age, sex, chronic inflammation, or anti-inflammatory medication can influence basal and post-radiation exposure gene expression. Recently, we used two transgenic mouse models that mimic chronic inflammation (*IL-10* knockout) [Mukherjee et al., 2019b] and anti-inflammation (dominant negative *p38α*^*MAPK*^) [Broustas et al., 2022]. We showed that generation of inflammatory conditions modified the expression of many radiation-responsive genes. However, we demonstrated that the inflammation status of mice has little impact on radiation dose classification. In contrast, we showed that age could have a profound impact on gene expression and diminish the predictive power of a previously created radiation biodosimetry gene list [Broustas et al., 2021]. Thus, previously identified radiation biomarker candidates may require adjustment to translate across age or sex and, thus, age- and sex-specific validation is crucial. In this article we examined the impact of sex alone or in combination with age on radiation dose reconstruction. Comparing differentially expressed genes between male and female irradiated mice, we found that young animals shared 26.4% of the total differentially expressed genes in either sex. In contrast, older male and female animals exposed to the same radiation dose shared only 14.5% of the differentially expressed genes. Of the initial radiation-responsive genes identified in young female and male animals, less than a third were also differentially expressed in older mice, particularly female animals, due to an increased replicate variability as visualized in a series of gene expression heatmaps (Fig. 2c). These results support the idea that aging is characterized by increased inter-individual variability of gene expression and are consistent with a less uniform radiation response of older mice compared with young animals [Patterson et al, 2022]. The combined effect of age and sex on radiation-responsive genes common to all age and sex mouse groups was also examined plotting fold-change of expression of each group against fold-change of expression of young male mice. Results revealed that fold-changes in young female mice were similar to those of young male mice, whereas fold-changes in old female were closer to old male mice, although sex differences in older animals were still significant. Thus, although sex does not appear to have an impact on the magnitude of up- or down-regulation of radiation-responsive gene expression in younger mice, it becomes a confounding factor in older age.

We used a novel approach to quantitatively reconstruct radiation dose, initially developed using human blood cells [Ghandhi et al., 2019], based on the gene expression data. The dose reconstruction algorithm was trained using the young male dataset and tested on the young female and old male and female datasets. Using this method, we were able to define a gene expression signature that could reconstruct the radiation dose of 4 Gy with root mean square error of ± 0.75 Gy with a coefficient of determination (R^2^) of 0.90. Of the groups tested, older female mice showed the greatest divergence of the estimated dose from the true dose (4 Gy), although the difference was not statistically significant. In the past, it has been demonstrated that classifiers derived using data from *ex vivo* irradiation of blood samples from a number of male and female human donors performed equally well over a range of radiation exposure doses (0.1 to 2 Gy) and was not affected by gender differences [Paul and Amundson, 2011]. However, it has been shown that sex differences may impact the accuracy of gene expression signatures to differentiate between low (e.g., 0.5 Gy) and control unirradiated mice, whereas at higher doses (e.g., 1-2 Gy), sex is not a confounding factor [Meadows et al., 2008]. Thus, the impact of sex on radiation biodosimetry may depend not only on age, but also on radiation dose. Considering this information, it will be necessary to test the robustness of our gene expression signature over a range of radiation doses, especially lower doses.

Gene expression profiling in radiation biodosimetry is not only useful for reconstructing physical absorbed radiation dose, but to predict the biological effect of the exposure and, thus, predict severity of injury [Port et al., 2016], as well as provide actionable genes that could be targeted to counteract the effects of radiation. We performed pathway analyses to identify canonical pathways and diseases or functions that were enriched in response to radiation in female young and old mice and compared with data from our previous study with male young and old animals [Broustas et al., 2021]. Cell death and specifically apoptosis plays multiple critical roles in the immune system, such as the negative selection of thymocytes and lymphocytes, as a defense mechanism against autoimmunity, and in the maintenance of proliferative homeostasis [Salminen et al., 2011]. It is well known that proliferating hematopoietic system cells predominantly undergo apoptosis in response to irradiation [Sarosiek et al., 2017]. Our analysis showed that the gene expression profile in peripheral blood cells from young mice of both sexes exposed to radiation correlated with an upregulation of apoptosis-related functions, whereas functions associated with quantity of many immune cell subtypes were markedly underrepresented. In contrast, cell death- or apoptosis-related functions were not enriched in irradiated old mice. Radiation-induced apoptosis of blood cells in young mice could subsequently be cleared by phagocytosis leading to inflammatory resolution and restoration of immune system function, whereas decreased apoptosis in response to irradiation could lead to induction and persistence of inflammation.

Phagocytosis is an indispensable cellular and molecular function tasked to maintain tissue homeostasis and innate immune balance [Li, 2013]. Phagocytosis clears apoptotic cells, as well as invading pathogens, and is mainly carried out by the so-called professional phagocytes that include macrophages, dendritic cells, and neutrophils [Li et al., 2013]. It is well established that females from various species, including humans and mice, have a more robust phagocytic activity than males and the phagocytic activity of neutrophils and macrophages is higher in females than males [Spitzer, 1999]. Although the role of sex hormones in phagocytosis has not been studied systematically, there is evidence that the phagocytic activity of macrophages is lower in males than females owing to the suppressive effects of androgens on male macrophage activity [Mondal and Rai, 1999]. The potential role of phagocytosis in radiation-induced processes is supported by our data that show phagocytosis-related biofunctions were upregulated in young irradiated female mice and these pathways were common between female and male mice following irradiation. In support of the previous studies that female mice have elevated phagocytic function, our results predict that unlike old male mice that are not enriched in phagocytosis-related functions, older female mice are enriched in some phagocytosis functions, albeit with smaller z-scores and p-values compared with young animals. Interestingly, few of these pathways were enriched in unirradiated mice. However, to confirm these observations, functional phagocytosis assays will be required.

Neutrophils are the predominant leukocyte subset in the human peripheral blood and play a major role in defense against microbial pathogens through their phagocytic activity and are essential for effective innate immunity [Mantovani et al., 2011]. They are recruited to sites of infection where they ingest and kill invading pathogens. Neutrophils are also required for the production of early, innate immune cell-derived IFNγ (probably by NK cells) and modulate the immune response and also contribute to ongoing inflammation in numerous diseases. Accordingly, neutropenia or neutrophil dysfunction leads to recurrent infections and life-threatening conditions [Mantovani et al., 2011; Jaillon et al., 2013]. Neutrophils are acting as phagocytes by releasing lytic enzymes from their granules and produce reactive oxygen species with antimicrobial potential [Mantovani et al., 2011]. One of the hallmarks associated with the antimicrobial and inflammatory action of neutrophils is the activation of a powerful oxidative burst, during which large amounts of oxygen are consumed and converted to superoxide radicals, which then act as precursor of hydrogen peroxide and other reactive oxygen species that are generated by their heme enzyme myeloperoxidase [Winterbourn et al., 2016]. Although we show that geriatric mice possess higher percentages of neutrophils both at steady state, and in the case of female mice after irradiation, as well, they exhibit a higher frequency of infection-related morbidities compared with young populations [Patterson et al., 2022], implying they may be dysfunctional. Furthermore, sex differences in neutrophil activity have been manifested with female hormones delaying apoptosis, as well as stimulating chemotaxis and recruitment of neutrophils to sites of infection [Jaillon et al., 2019]. In addition, in humans, sex hormones modulate the production of reactive oxygen species intermediate by neutrophils [Bouman et al., 2005]. However, although neutrophil activation protects the organism from infections, their activation releases cytotoxic mediators that can cause tissue damage, such as myocardial infarction and autoimmune disease [Luster et al., 2005]. While young female mice are predicted to have a more robust inflammatory response immediately following radiation exposure than young male or old animals, it would be necessary to perform longitudinal studies to determine the time it takes for neutrophil activity to return to basal levels following radiation exposure and whether they contribute to the resolution or propagation of inflammation response to irradiation and thus whether the play a protective or harmful role in tissue and organismal injury. In general, myeloid cell related functions, which are the first line of defense to infection after an acute radiation exposure were significantly overrepresented in young female and male mice, less represented in older female mice and absent in older male mice, despite the well-established age-biased increased myeloid cell production in both humans and animals [Elias et al., 2017], but consistent with their relatively impaired activity.

The long-term goal of our studies is to develop gene expression signatures robust enough to detect radiation exposure, irrespective of confounding factors, such as age, sex, or co-morbidities. In this and a previous publication [Broustas et al., 2021], we studied the impact of age and sex on the accuracy of a gene expression signature to reconstruct the correct radiation dose. Our data show that although age and sex can lessen the robustness of the reconstructed dose estimate, deviations are not statistically significant. However, although age and sex appear to have limited influence on radiation dose estimates, at least at 4 Gy X-rays, gene ontology analysis demonstrated that there are significant variations among enriched biological pathways, diseases, and functions among female and male animals combined with young and old age and, thus, sex may have an impact on the severity of radiation injury. For this reason, it is crucial to include both male and female individuals in the study of radiation biodosimetry. However, to validate the findings of the present study, functional studies will be necessary to causally connect the differentially enriched pathways with severity of radiation damage.

## Additional Information

Supplementary information is available for this paper.

## Acknowledgements

Analyses were performed using BRB-ArrayTools developed by Dr. Richard Simon and BRB-ArrayTools Development Team.

## Statement of Ethics

All animal experiments were conducted in accordance with applicable federal and state guidelines based on approved by the Columbia University Institutional Animal Care and Use Committee (approval number AC-AAAT6450).

## Competing interests

The authors declare no competing interests.

## Funding Sources

This work was supported by a pilot grant from the Center for High-Throughput Minimally-Invasive Radiation Biodosimetry, National Institute of Allergy and Infectious Diseases Grant number U19AI067773.

## Author contributions

C.G.B. designed the study, performed the experiments and analyzed the primary data, interpreted the results and wrote the manuscript. A.J.D. assisted conducting the animal experiments. I.S. performed the dose reconstruction analysis. S.A.A. contributed to data analysis and interpretation, and writing the manuscript. All authors read and approved the final manuscript.

## Data Availability Statement

The microarray data for the female mice generated in this study have been deposited in the National Center for Biotechnology Information Gene Expression Omnibus (GEO) database with accession number GSE133451 (https://www.ncbi.nlm.nih.gov/geo/query/acc.cgi?acc=GSE133451). The microarray data for the male mice study has been deposited in GEO under the accession number GSE132559.

**Supplementary Fig. 1 (related to Fig. 2).**
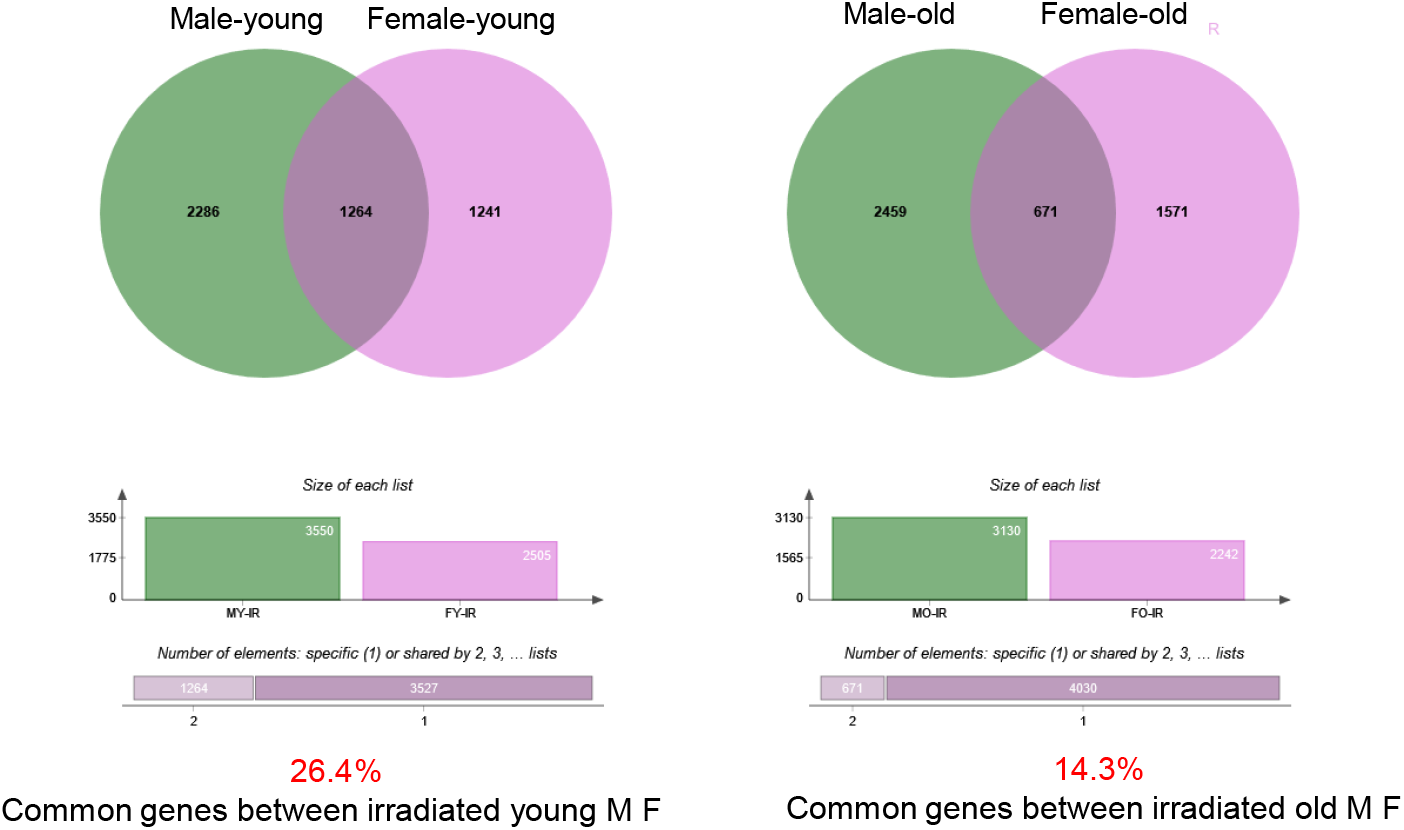
Common differentially expressed genes between irradiated young male and irradiated young female mice (*left*) and irradiated old male and irradiated old female mice (*right*). Below the Venn diagrams, the total number of unique and common differentially expressed genes in both sexes is depicted and the percentage of common genes is calculated. M: male; F: female.

**Table S1a.** Immune cell death-related functions between young and old female irradiated mice.

|  | Young | Old |
| --- | --- | --- |
| Diseases and Functions Annotation | z-score |  |
| Cell death of immune cells | 3.920 | 0.604 |
| Cell death of blood cells | 3.774 | 0.187 |
| Cell death of B lymphocytes | 2.706 | 0.000 |
| Apoptosis of leukocytes | 2.705 | -0.449 |
| Cell death of myeloid cells | 2.537 | 0.000 |
| Cell death of lymphocytes | 2.370 | 0.000 |
| Cell death of lymphatic system cells | 2.198 | 0.000 |
| Cell death of lymphoid cells | 2.182 | 0.000 |
| Cell death of mononuclear leukocytes | 2.168 | -0.033 |
| Production of antibody | -2.346 | -2.020 |
| Quantity of immunoglobulin | -2.659 | -2.204 |
| Quantity of IgG1 | -3.046 | 0.000 |
| Quantity of lymphoid tissue | -2.378 | -0.637 |
| Quantity of B lymphocytes | -3.155 | 0.000 |
| Quantity of IgG | -2.562 | 0.000 |

**Table S1b.** Immune cell death-related functions between young and old female and male irradiated mice.

|  | Female |  | Male |  |
| --- | --- | --- | --- | --- |
|  | 2-mo | 24-mo | 2-mo | 21-mo |
| Diseases and Functions Annotation | z-score |  |  |  |
| Cell death of immune cells | 3.920 | 0.604 | 2.285 | -0.276 |
| Cell death of blood cells | 3.774 | 0.187 | 2.152 | -0.271 |
| Cell death of B lymphocytes | 2.706 | 0.000 | 2.973 | 1.646 |
| Apoptosis of leukocytes | 2.705 | -0.449 | 2.700 | 0.411 |
| Cell death of myeloid cells | 2.537 | 0.000 | 0.000 | 0.000 |
| Cell death of lymphocytes | 2.370 | 0.000 | 3.063 | 1.051 |
| Cell death of lymphatic system cells | 2.198 | 0.000 | 2.774 | 1.131 |
| Apoptyosis of lymphoid cells | 2.056 | 0.000 | 2.620 | 0.000 |
| Apoptosis of mononuclear leukocytes | 2.008 | -0.033 | 2.632 | 0.697 |
| Quantity of lymphoid tissue | -2.378 | -0.637 | -3.742 | -2.028 |
| Quantity of B lymphocytes | -3.155 | 0.000 | -4.166 | -3.120 |
| Quantity of T lymphocytes | 0.000 | 0.000 | -2.538 | -1.260 |
| Quantity of lymphocytes | 0.000 | 0.000 | -3.206 | -1.960 |

**Table S2a.** Phagocytosis-related functions between young and old female mice exposed to 4 Gy x-rays.

|  | Young | Old |
| --- | --- | --- |
| Diseases and Functions Annotations | z-score |  |
| Endocytosis | 5.861 | 3.029 |
| Cell movement of phagocytes | 5.113 | 3.382 |
| Engulfment of cells | 5.100 | 3.082 |
| Endocytosis by eukaryotic cells | 5.068 | 2.496 |
| Phagocytosis | 5.061 | 2.470 |
| Migration of phagocytes | 4.519 | 2.150 |
| Phagocytosis of cells | 4.467 | 2.643 |
| Chemotaxis of phagocytes | 4.317 |  |
| Recruitment of phagocytes | 4.266 | 3.052 |
| Activation of phagocytes | 3.958 |  |
| Phagocytosis of myeloid cells | 3.678 |  |
| Engulfment of blood cells | 3.595 |  |
| Engulfment of leukocytes | 3.507 |  |
| Phagocytosis of blood cells | 3.506 |  |
| Phagocytosis of leukocytes | 3.415 |  |
| Phagocytosis of phagocytes | 3.269 |  |
| Engulfment of phagocytes | 2.971 |  |
| Degranulation of phagocytes | 2.69 |  |
| Cell death of phagocytes | 2.603 |  |
| Cellular infiltration by phagocytes | 2.497 |  |
| Immune response of phagocytes | 2.325 |  |
| Cell viability of phagocytes | 2.259 |  |
| Apoptosis of phagocytes | 2.073 |  |
| Binding of professional phagocytic cells |  | 2.305 |

**Table S2b.** Comparison of phagocytosis-related functions between male and female mice exposed to 4 Gy x-rays.

|  | Young | Old |
| --- | --- | --- |
| Diseases or Functions Annotation | z-score |  |
| Phagocytosis | 4.372 |  |
| Phagocytosis of cells | 4.123 | 2.042 |
| Engulfment of cells | 3.943 |  |
| Internalization of cells | 3.778 |  |
| Cell movement of phagocytes | 3.712 |  |
| Endocytosis | 3.519 |  |
| Endocytosis by eukaryotic cells | 3.416 | 2.162 |
| Cell death of phagocytes | 3.163 |  |
| Phagocytosis of blood cells | 3.145 |  |
| Phagocytosis of phagocytes | 3.127 |  |
| Apoptosis of phagocytes | 3.054 |  |
| Phagocytosis of myeloid cells | 3.020 |  |
| Phagocytosis of leukocytes | 3.008 |  |
| Engulfment of myeloid cells | 2.925 |  |
| Migration of phagocytes | 2.852 |  |
| Engulfment of blood cells | 2.790 |  |
| Phagocytosis of antigen presenting cells | 2.700 |  |
| Engulfment of leukocytes | 2.648 |  |
| Engulfment of phagocytes | 2.548 |  |
| Engulfment by macrophages | 2.236 |  |
| Recruitment of phagocytes | 2.112 |  |

**Table S3a.** Neutrophil-associated functions among irradiated mice of different sex and age.

|  | female |  | male |  |
| --- | --- | --- | --- | --- |
|  | 2-mo | 24-mo | 2-mo | 21-mo |
| Diseases or Functions Annotation | z-score |  |  |  |
| Recruitment of neutrophils | 3.829 |  | 2.944 |  |
| Cell movement of neutrophils | 3.096 | 3.280 |  |  |
| Chemotaxis of neutrophils | 4.523 |  |  |  |
| Migration of neutrophils | 2.700 |  |  |  |
| Infiltration by neutrophils | 2.200 |  |  |  |
| Degranulation of neutrophils | 2.121 |  |  |  |

**Table S3b.**
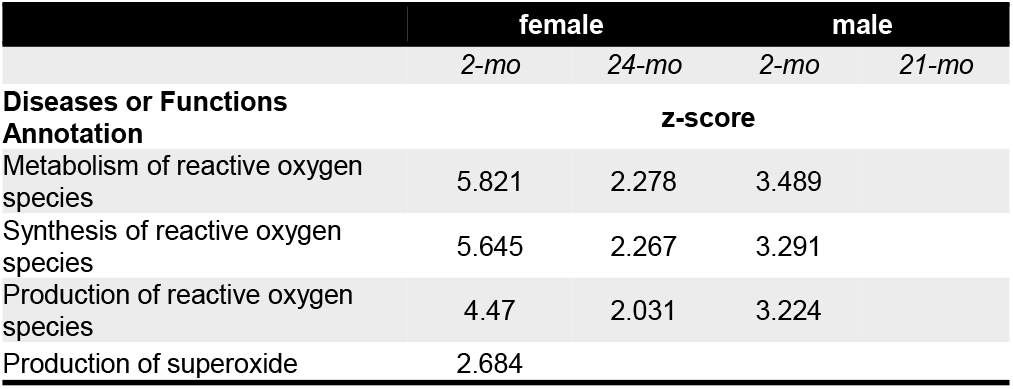
Reactive oxygen species synthesis and metabolism-associated functions among irradiated mice of different sex and age.

**Table S3c.** Myeloid-associated functions among irradiated mice of different sex and age.

|  | female |  | male |  |
| --- | --- | --- | --- | --- |
|  | 2-mo | 24-mo | 2-mo | 21-mo |
| Diseases or Functions Annotation | z-score |  |  |  |
| Cell movement of myeloid cells | 5.22 | 3.508 | 2.454 |  |
| Recruitment of myeloid cells | 4.338 |  |  |  |
| Activation of myeloid cells | 4.184 |  | 3.665 |  |
| Chemotaxis of myeloid cells | 4.523 | 3.315 |  |  |
| Binding of myeloid cells | 3.835 | 2.975 | 3.218 |  |
| Phagocytosis of myeloid cells | 3.678 |  |  |  |
| Adhesion of myeloid cells | 3.386 |  |  |  |
| Migration of myeloid cells | 3.311 |  |  |  |
| Response of myeloid cells | 3.155 |  | 3.089 |  |
| Cellular infiltration by myeloid cells | 2.885 |  |  |  |
| Engulfment of myeloid cells |  |  | 3.745 |  |
| Recruitment of myeloid cells |  |  | 2.713 |  |

## Notes

### Competing Interest Statement

The authors have declared no competing interest.

## References

Amundson SA. The transcriptomic revolution and radiation biology. Int J Radiat Biol. 2022;98(3):428–438.

Bakker OB, Aguirre-Gamboa R, Sanna S, Oosting M, Smeekens SP, Jaeger M, et al. Integration of multi-omics data and deep phenotyping enables prediction of cytokine responses. Nat Immunol. 2018;19(7):776–786.

Bardou P, Mariette J, Escudié F, Djemiel C, Klopp C. jvenn: an interactive Venn diagram viewer. BMC Bioinformatics. 2014;15(1):293.

Boissier J, Chlichlia K, Digon Y, Ruppel A, Moné H. Preliminary study on sex-related inflammatory reactions in mice infected with Schistosoma mansoni. Parasitol Res. 2003;91(2):144–50.

Bouman A, Heineman MJ, Faas MM. Sex hormones and the immune response in humans. Hum Reprod Update. 2005;11(4):411–23.

Broustas CG, Mukherjee S, Pannkuk EL, Laiakis EC, Fornace AJ, Amundson SA. Effect of the p38 Mitogen-Activated Protein Kinase Signaling Cascade on Radiation Biodosimetry. Radiat Res. 2022;198(1):18–27.

Broustas CG, Duval AJ, Amundson SA. Impact of aging on gene expression response to X-ray irradiation using mouse blood. Sci Rep. 2021;11(1):10177.

Broustas CG, Harken AD, Garty G, Amundson SA. Identification of differentially expressed genes and pathways in mice exposed to mixed field neutron/photon radiation. BMC Genomics. 2018;19(1):504.

Broustas CG, Xu Y, Harken AD, Garty G, Amundson SA. Comparison of gene expression response to neutron and X-ray irradiation using mouse blood. BMC Genomics. 2017a;18(1):2.

Broustas CG, Xu Y, Harken AD, Chowdhury M, Garty G, Amundson SA. Impact of Neutron Exposure on Global Gene Expression in a Human Peripheral Blood Model. Radiat Res. 2017b;187(4):433–440.

Dressman HK, Muramoto GG, Chao NJ, Meadows S, Marshall D, Ginsburg GS, et al. Gene expression signatures that predict radiation exposure in mice and humans. PLoS Med. 2007;4(4):e106.

Elias HK, Bryder D, Park CY. Molecular mechanisms underlying lineage bias in aging hematopoiesis. Semin Hematol. 2017;54(1):4–11.

Fulop T, Larbi A, Dupuis G, Le Page A, Frost EH, Cohen AA, Witkowski JM, Franceschi C. Immunosenescence and Inflamm-Aging As Two Sides of the Same Coin: Friends or Foes? Front Immunol. 2018;8:1960.

Furman D, Hejblum BP, Simon N, Jojic V, Dekker CL, Thiébaut R, et al. Systems analysis of sex differences reveals an immunosuppressive role for testosterone in the response to influenza vaccination. Proc Natl Acad Sci U S A. 2014;111(2):869–74.

Ghandhi SA, Shuryak I, Morton SR, Amundson SA, Brenner DJ. New Approaches for Quantitative Reconstruction of Radiation Dose in Human Blood Cells. Sci Rep. 2019;9(1):18441.

Goronzy JJ, Li G, Yang Z, Weyand CM. The janus head of T cell aging - autoimmunity and immunodeficiency. Front Immunol. 2013;4:131.

Giefing-Kröll C, Berger P, Lepperdinger G, Grubeck-Loebenstein B. How sex and age affect immune responses, susceptibility to infections, and response to vaccination. Aging Cell. 2015;14(3):309–21.

Hernández L, Terradas M, Camps J, Martín M, Tusell L, Genescà A. Aging and radiation: bad companions. Aging Cell. 2015;14(2):153–61.

Hochberg Y, Benjamini Y. More powerful procedures for multiple significance testing. Stat Med. 1990;9, 811–8.

Jaillon S, Berthenet K, Garlanda C. Sexual Dimorphism in Innate Immunity. Clin Rev Allergy Immunol. 2019;56(3):308–321.

Jaillon S, Galdiero MR, Del Prete D, Cassatella MA, Garlanda C, Mantovani A. Neutrophils in innate and adaptive immunity. Semin Immunopathol. 2013;35(4):377–94.

Klein SL, Flanagan KL. Sex differences in immune responses. Nat Rev Immunol. 2016;16(10):626–38.

Li S, Lu X, Feng JB, Tian M, Wang J, Chen H, et al. Developing Gender-Specific Gene Expression Biodosimetry Using a Panel of Radiation-Responsive Genes for Determining Radiation Dose in Human Peripheral Blood. Radiat Res. 2019;192(4):399–409.

Li W. Phagocyte dysfunction, tissue aging and degeneration. Ageing Res Rev. 2013;12(4):1005–12.

Luster AD, Alon R, von Andrian UH. Immune cell migration in inflammation: present and future therapeutic targets. Nat Immunol. 2005;6(12):1182–90.

Mantovani A, Cassatella MA, Costantini C, Jaillon S. Neutrophils in the activation and regulation of innate and adaptive immunity. Nat Rev Immunol. 2011;11(8):519–31.

Márquez EJ, Chung CH, Marches R, Rossi RJ, Nehar-Belaid D, Eroglu A, Mellert DJ, Kuchel GA, Banchereau J, Ucar D. Sexual-dimorphism in human immune system aging. Nat Commun. 2020;11(1):751.

Meadows SK, Dressman HK, Muramoto GG, Himburg H, Salter A, Wei Z, et al. Gene expression signatures of radiation response are specific, durable and accurate in mice and humans. PLoS One. 2008;3(4):e1912.

Melgert BN, Oriss TB, Qi Z, Dixon-McCarthy B, Geerlings M, Hylkema MN, Ray A. Macrophages: regulators of sex differences in asthma? Am J Respir Cell Mol Biol. 2010;42(5):595–603.

Mondal S, Rai U. Sexual dimorphism in phagocytic activity of wall lizard’s splenic macrophages and its control by sex steroids. Gen Comp Endocrinol. 1999;116(2):291–8.

Mukherjee S, Grilj V, Broustas CG, Ghandhi SA, Harken AD, Garty G, Amundson SA. Human Transcriptomic Response to Mixed Neutron-Photon Exposures Relevant to an Improvised Nuclear Device. Radiat Res. 2019a;192(2):189–199.

Mukherjee S, Laiakis EC, Fornace AJ Jr, Amundson SA. Impact of inflammatory signaling on radiation biodosimetry: mouse model of inflammatory bowel disease. BMC Genomics. 2019b;20(1):329.

Patterson AM, Vemula S, Plett PA, Sampson CH, Chua HL, Fisher A, et al. Age and Sex Divergence in Hematopoietic Radiosensitivity in Aged Mouse Models of the Hematopoietic Acute Radiation Syndrome. Radiat Res. 2022; doi: 10.1667/RADE-22-00071.1.

Paul S, Amundson SA. Gene expression signatures of radiation exposure in peripheral white blood cells of smokers and non-smokers. Int J Radiat Biol. 2011;87(8):791–801.

Peters MJ, Joehanes R, Pilling LC, Schurmann C, Conneely KN, Powell J, et al. The transcriptional landscape of age in human peripheral blood. Nat Commun. 2015;6:8570.

Piasecka B, Duffy D, Urrutia A, Quach H, Patin E, Posseme C, et al. Distinctive roles of age, sex, and genetics in shaping transcriptional variation of human immune responses to microbial challenges. Proc Natl Acad Sci U S A. 2018;115(3):E488–E497.

Port M, Herodin F, Valente M, Drouet M, Lamkowski A, Majewski M, et al. First Generation Gene Expression Signature for Early Prediction of Late Occurring Hematological Acute Radiation Syndrome in Baboons. Radiat Res. 2016;186(1):39–54.

Salminen A, Ojala J, Kaarniranta K. Apoptosis and aging: increased resistance to apoptosis enhances the aging process. Cell Mol Life Sci. 2011;68(6):1021–31.

Sarosiek KA, Fraser C, Muthalagu N, Bhola PD, Chang W, McBrayer SK, et al. Developmental Regulation of Mitochondrial Apoptosis by c-Myc Governs Age- and Tissue-Specific Sensitivity to Cancer Therapeutics. Cancer Cell. 2017;31(1):142–156.

Spitzer JA. Gender differences in some host defense mechanisms. Lupus. 1999;8(5):380–3.

vom Steeg LG, Klein SL. SeXX Matters in Infectious Disease Pathogenesis. PLoS Pathog. 2016;12(2):e1005374.

Whitney AR, Diehn M, Popper SJ, Alizadeh AA, Boldrick JC, Relman DA, et al. Individuality and variation in gene expression patterns in human blood. Proc Natl Acad Sci U S A. 2003;100(4):1896–901.

Winterbourn CC, Kettle AJ, Hampton MB. Reactive Oxygen Species and Neutrophil Function. Annu Rev Biochem. 2016;85:765–92.

Wright GW, Simon RM. A random variance model for detection of differential gene expression in small microarray experiments. Bioinformatics. 2013;19, 2448–55.

Xia HJ, Zhang GH, Wang RR, Zheng YT. The influence of age and sex on the cell counts of peripheral blood leukocyte subpopulations in Chinese rhesus macaques. Cell Mol Immunol. 2009;6(6):433–40.

